# Conformational dynamics of the HIV Vif protein complex

**DOI:** 10.1101/483099

**Authors:** K. A. Ball, L. M. Chan, D. J. Stanley, E. Tierney, S. Thapa, H. M. Ta, L. A. Burton, J. M. Binning, M. P. Jacobson, J. D. Gross

## Abstract

HIV-1 viral infectivity factor (Vif) is an intrinsically disordered protein responsible for the ubiquitination of the APOBEC3 antiviral proteins. Vif folds when it binds the Cullin-RING E3 ligase CRL5 and the transcription cofactor CBF-β. A five-protein complex containing the substrate receptor (Vif, CBF-β, Elongin-B, Elongin-C) and Cullin5 (CUL5) has a published crystal structure, but dynamics of this VCBC-CUL5 complex have not been characterized. Here, we use Molecular Dynamics (MD) simulations and NMR to characterize the dynamics of the VCBC complex with and without CUL5 and APOBEC3 bound. Our simulations show that the VCBC complex undergoes global dynamics involving twisting and clamshell opening of the complex, while VCBC-CUL5 maintains a more static conformation, similar to the crystal structure. This observation from MD is supported by methyl-transverse relaxation optimized spectroscopy (methyl-TROSY) NMR data, which indicates that the entire VCBC complex without CUL5 is dynamic on the μs-ms timescale. Vif binds APOBEC3 to recruit it to the complex, and methyl-TROSY NMR shows that the VCBC complex is more conformationally restricted when bound to APOBEC3F, consistent with our MD simulations. Vif contains a flexible linker region located at the hinge of the VCBC complex, which changes conformation in conjuction with the global dynamics of the complex. Like other ubiquitin substrate receptors, VCBC can exist alone or in complex with CUL5 in cells. Accordingly, the VCBC complex could be a good target for therapeutics that would inhibit full assembly of the ubiquitination complex by stabilizing an alternate VCBC conformation.

## Introduction

Human restriction factors provide a critical line of defense against viral pathogens. However, many viruses encode proteins to counteract restriction factors and allow for persistent infection in host. A prime example of this host-pathogen conflict is provided by the HIV-1 protein-viral infectivity factor (Vif), which targets the host antiviral APOBEC3 proteins. In the absence of Vif, APOBEC3 family members are encapsidated into HIV virions and inhibit viral replication primarily by deamination of cytidines to uridines in the viral cDNA during reverse transcription (1). The resulting hypermutation renders the viral infection nonproductive (2). Nearly all lentiviruses encode a Vif protein that counteracts the APOBEC3 family members, targeting them for destruction by ubiquitin mediated proteolysis (3). Vif directly binds to APOBEC3s and hijacks a Cullin-RING E3 ligase (CRL), which acts in the last step of a three-enzyme E1-E2-E3 cascade. CRLs consist of a substrate receptor, a Cullin backbone and a RING-box protein (RBX), which activates and delivers a ubiquitin carrier protein (E2). HIV-1 Vif recruits APOBEC3 family members to CRL5, which contains the adaptor proteins Elongin B (EloB) and Elongin C (EloC) that connect the substrate-binding protein to Cullin-5 (CUL5). For example, in uninfected cells, Suppressor Of Cytokine Signaling (SOCS) proteins act as substrate binding proteins that promote interaction of the EloB and EloC proteins with CUL5 in order to attenuate cytokine signaling by ubiquitination (Fig. 1a). As shown in Fig. 1b, Vif hijacks CRL5 by replacing SOCS2 as the substrate-binding protein, interacting directly with EloC and CUL5 (4). Vif also recruits core binding factor subunit-β (CBF-β), a transcription cofactor, to the CRL5 complex (5, 6). Though not present in the cellular E3 ligase complex, CBF-β is required for the formation of a functional HIV-1 Vif E3 holoenzyme both in vitro and during HIV infection (5, 6).

**Figure 1.**
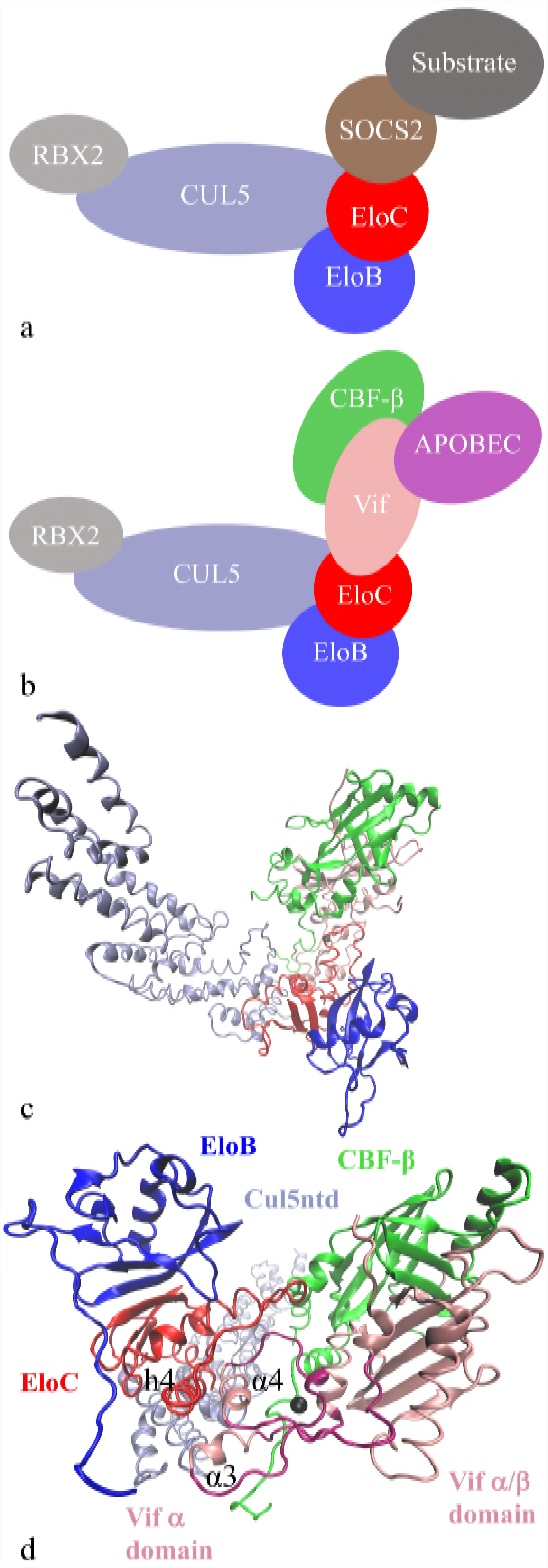
Schematic diagram of (a) the host SOCS-CLR5 E3 ubiquitin ligase complex and (b) the HIV-Vif E3 ubiquitin ligase complexes. The VCBC-CUL5ntd crystal structure (PDB: 4N9F) is shown from the same orientation as the diagram (c) and rotated (d) to more clearly show the two Vif domains connected by the Vif linker region (dark pink) containing the zinc ion shown as a black sphere. Vif helices α3 and α4 and EloC helix h4 are labeled.

Vif is an intrinsically disordered protein (7), yet Guo *et al.* solved a crystal structure of a complex between Vif, CBF-β, EloB, EloC, (termed VCBC) and the N-terminal domain of CUL5 (CUL5ntd) (4). This structure revealed that the four-protein VCBC is a co-folded unit, and that Vif is folded into a larger N-terminal α/β-domain and smaller C-terminal α-domain that are connected through a linker region containing a zinc binding motif (4). For this APOBEC3 substrate receptor complex to form, Vif must adopt a folded conformation, stabilized by interactions with CBF-β and EloBC (6). The α-domain of Vif binds ELOBC and CUL5 by mimicking the SOCS2 interaction surface. It contains a BC-box motif required to interact with the EloB and EloC heterodimer (EloBC) and a surface that directly binds CUL5 (Fig. 1d). The α/β-domain of Vif is responsible for interacting with CBF-β, and through this interaction, CBF-β stabilizes the two loosely packed helices in the α/β-domain of Vif and forms an extended intermolecular β-sheet with Vif’s N-terminus (4–6). Between these two domains, whose secondary structure is stabilized by their interacting partners, the Vif linker region is less structured, stabilized by a zinc ion that interacts with the HCCH arrangement of His and Cys residues in the Vif linker region (4). We hypothesized that this unstructured linker region, located in the center point of the VCBC structure, may form a hinge that determines flexibility and conformational changes of the complex. Although the intrinsically disordered Vif protein folds as part of the CRL complex, flexibility retained in the linker region may be critical to allowing different regions of the complex to reorient, allowing the ubiquitination transfer from RBX to A3 to occur.

CUL5 is present in the Guo *et al.* crystal structure, but it is not required for Vif to form the stable VCBC complex (6). Rather, CUL5 stabilizes the already formed VCBC complex when it binds, interacting directly with EloC and with Vif (4). The Vif HCCH zinc-binding motif is also required for CUL5 binding, though it does not directly interact with CUL5 (4). As noted by Guo and Zhou, sometimes when a protein partner, like CUL5 is removed, favorable interactions are lost, causing proteins to sample more states and show an increase in conformational entropy (8). Furthermore, a multitude of substrate receptors must share a given CRL backbone, which is promoted by cycles of NEDD8 conjugation, deconjugation and substrate receptor exchange (9). There are also at least three distinct A3 binding interfaces on Vif: the A3F-like binding site (A3F/C/D), the A3G-binding site, and the A3H-binding site (10, 11), which may have different binding affinities for different conformations of the VCBC complex. Accordingly, understanding the structural dynamics of the VCBC subunit not bound to the CUL5 scaffold could be advantageous for understanding the complete process of A3 ubiquitination by Vif. Additionally, full assembly of the CRL complex potentially could be inhibited if alternate conformations of the VCBC complex incompatible with CUL5 binding are stabilized, e.g., by small molecule binding.

Previous molecular dynamics (MD) simulation studies by Liu and Nussinov have shown that ubiquitination complexes are not completely rigid (12–14). In their simulations of several CRL substrate binding proteins and Cullins 1 and 5, they found that conformations different from those in the crystal structure are sampled. Specifically, the substrate binding protein that links the adaptor (in our case EloB-EloC) to the substrate (in our case APOBEC3) was found to contain a flexible linker region that allowed the substrate-binding domain to reorient relative to the BC-box domain (12, 13). Liu and Nussinov also found that this conformational flexibility was allosterically regulated by binding of both the adaptor and substrate proteins, sometimes restricting the conformations sampled (12, 13). In the VCBC complex, Vif plays the role of the substrate binding protein, with the help of CBF-β, and also contains a linker region between its two domains. No structures have been solved with Vif bound to a substrate A3 protein, but several mutational studies have been performed that indicate the interface on the α/β domain of Vif is where the A3s bind (15, 16). Since the Vif-ubiquitination complex involves the large-scale transfer of the ubiquitin moiety from the E2 at one end of the complex to the substrate at the other, alternate conformations could be important for function of the complex (12). In this study, we compare the conformational ensembles and alternate states of the HIV-1 Vif complex and investigate the role Vif plays in the flexibility of the complex.

We used MD simulations to investigate conformational dynamics of the VCBC complex and how CUL5 binding modulates the conformational ensemble. Simulations enable direct comparison of conformational states for the VCBC complex with and without CUL5 bound, including states that may be difficult to capture experimentally because of their low occupancy. We found that the VCBC-CUL5 complex samples states similar to that found in the crystal structure, while the VCBC complex undergoes large-scale conformational changes involving the flexible hinge region of Vif. MD simulations of VCBC-A3F based on a modeled interface (17) also indicated that the VCBC complex is conformationally restricted when bound to the C-terminal domain of A3F (A3Fctd). We also carried out methyl-transverse relaxation optimized spectroscopy (TROSY) NMR experiments, which support the existence of multiple alternate conformations of the VCBC complex that interconvert on the μs-ms timescale, causing extensive peak broadening; however, the VCBC complex bound to A3Fctd displays reduced VCBC dynamics. Thus, both computational simulations and NMR experiments demonstrate that the HIV-1 Vif substrate receptor (VCBC) is a dynamic complex that samples multiple conformations, and these dynamics are partially quenched upon binding to substrate or additional E3 ligase components. Structural adaptability in Vif is likely important for it to bind multiple A3 proteins through different binding modes and to carry out the catalytic step of ubiquitin transfer. By revealing alternate conformational states, simulations could also provide strategies to discover future therapeutics that inhibit A3 ubiquitination and degradation, resulting in A3 encapsidation and inhibition of viral replication.

## Materials and Methods

### Model building & system preparation

Crystal structures are available for VCBC-CUL5ntd and A3F-CTD. MD simulations were run on four different constructs of the HIV-Vif complexes: VCBC, VCBC-CUL5ntd, VCBC-A3Fctd, and VCBC with the shorter EloB102. The initial coordinates for each of these simulations came from the crystal structure of VCBC-CUL5ntd, PDB 4N9F (4). The crystal structure PDB file was missing coordinates for several unstructured regions of the proteins: Vif 1-2 and 173-176, CBF-β 1 and 157-165, EloB 80-82 and 99-102, EloC 50-57 and 112, CUL5 119-130. These were built using the PLOP homology modeling software to predict the conformation of the missing residues (18). Because the HIV-Vif consensus sequence was used for the NMR experiments (19), a homology model of the consensus Vif was built in Prime (20, 21). Histidine protonation states were optimized using Maestro (22). The crystal structure of VCBC-CUL5ntd contained a truncated EloB protein of only 102 residues, while the wild-type EloB has 118 residues. Therefore, for all simulated constructs except VCBC with EloB102, the starting structure for the C-terminus of EloB was obtained from a crystalized structure of the HIV Vif SOCS-box of EloBC, PDB 2MA9, which has the full length EloB (23). To combine the two structures, residues 1-78 of EloB were aligned in the two structures using Visual Molecular Dynamics (VMD) (24, 25), and residues 79-102 of EloB from 4N9F were replaced with residues 79-118 of EloB from 2MA9. For the VCBC and VCBC-A3Fctd constructs, the CUL5ntd atoms were removed.

For the simulations of VCBC-A3Fctd, the starting structure was obtained from the model by Richards *et al.* (26). To combine the Guo *et al.* VCBC-CUL5 crystal structure with the Richards *et al.* modeled structure of the A3F-Vif interface, the Vif interface residues (42–52 and 75-85) were aligned in the two structures using VMD (24), and the structure of A3Fctd from the Richards *et al.* model was added to the VCBC structure. Because the solubility enhanced A3Fctd construct used in our NMR experiments had 11 mutations (Y196D, H247G, C248R, C259A, F302K, W310D, Y314A, Q315A, K355D, K358D, F363D), our simulations were performed on this construct for comparative purposes. The VCBC-A3Fctd structure was changed to match this experimental sequence and the mutated side chains were modeled in with LEaP (27). Six missing residues at the N-terminus of the A3Fctd were modeled in using PLOP (21).

The Amber99SB protein force field was used for our MD simulations with the TIP3P water model (28, 29). The Amber 16 LEaP module was used to prepare each complex for simulation and create Amber parameter and input coordinate files. To model zinc ions and coordinated residues, we treated the zinc ion with a four-point charge representation as reported (30). The proteins were surrounded with at least 10 Å of water on all sides in a periodic box, and Na^+^ ions were added to neutralize the system (Table S1 in the Supporting Material).

### Molecular dynamics simulations

Energy minimization was performed for each complex prior to running MD simulations, initially with restraints of 500 kcal/(mol-Å^2^) on the protein atoms to allow the solvent to minimize, and then with no restraints to allow the protein structure to minimize away from any high-energy residue conformations. 500 steps of steepest descent minimization were performed followed by 1000 steps of gradient minimization for each round. The systems were then equilibrated for 20 ps at constant volume while raising the temperature from 0 K to 300 K, with restraints of 10 kcal/(mol-Å^2^) on protein atoms. Two more constant pressure equilibrations were performed to equilibrate the density of the systems with 1 kcal/(mol-Å^2^) restraints on the protein for 20 ps, and then without restrains for 1 ns. Dimensions of the periodic box after equilibration for each construct are given in the Table S1. All MD simulations were performed on GPUs using the cuda version of the *pmemd* module in the Amber package (27). The Andersen thermostat was used to hold the temperature constant at 300 K (31, 32), and the Berendsen barostat with isotropic position scaling and a relaxation time of 1 ps was used to hold the system at 1 bar (33). The particle-mesh Ewald procedure was used to handle long-range electrostatic interactions with a non-bonded cutoff of 9 Å. After the initial 1 ns equilibration, four independent production simulations were run for each construct starting from the same equilibrated structure, but with new initial, randomized, velocities. The total simulation time for each construct is given in Table S1. Snapshots were saved every 5 ps for the production simulations.

### Structure and trajectory analysis

Analysis of the molecular dynamics simulations were performed using the *cpptraj* module of Amber (27), and in-house python scripts. Interatomic distances, dihedral angles, root-mean-squared-deviations (RMSD) of protein structures, root-mean-squared fluctuations, and hydrogen bond percentage were all calculated using *cpptraj*. Cartesian Principal Component Analysis (PCA) was also calculated with *cpptraj*, based on the C_α_ position for each residue in the structured regions of the VCBC complex (excluding flexible tails and loops). The principal components (PCs) were taken as the first eigenvectors, *v*, of the covariance matrix, *M*, for these C_α_ positions. The data used to construct this covariance matrix included the ensemble of structures from both the VCBC simulations and the VCBC-CUL5 simulations. The data from each of the simulated constructs was then projected onto these same PCs to compare the global motions of the VCBC complex for each system.

All histograms were computed by combining the independent simulations for a given construct (snapshots spaced every 5 ps) and binning over this total ensemble of structures. To determine the timescale of the VCBC conformational exchange, the autocorrelation time, *τ*, was calculated for each metric. Because the autocorrelation time for the first two PCs was only one order of magnitude smaller than our total simulation time for each independent simulation, we treated the data from one simulation as correlated for purposes of statistical analysis. Only data from separate simulations were treated independently. All protein structure figures were created using Visual Molecular Dynamics (VMD) (24, 25). To cluster the VCBC ensemble by principal components 1 and 2, we used the agglomerative clustering method in the Python machine learning module, scikit-learn (34), which recursively merges pairs of clusters that minimally increase a given linkage distance. We used the Ward linkage, which minimizes the variance of the clusters being merged so that there will be greater variance in the data in different clusters than within one cluster. The number of clusters was set to 3 because we observed two very similar clusters when we increased the number to 4 clusters. To sort the VCBC-CUL5ntd data into clusters, we fit a linear support vector classification machine to the VCBC clusters with scikit-learn (34) and used that fit to predict the VCBC-CUL5 cluster membership.

### Assessment of simulation convergence

To make sure that we compared simulations that had equilibrated sufficiently away from the initial crystal structure, we divided each of the simulations into 100-ns blocks and performed PCA on the ensemble of structures from the four independent simulations for each block. Table S2 in the Supporting Material shows the first five eigenvalues for each block. We then took the sum of the first five eigenvalues, which corresponds to the variance captured by the first five principal components, and plotted how this quantity changes for each simulation block (Fig. S1 in the Supporting Material). We observed that for the VCBC-CUL5ntd and VCBC simulations, the variance increased and then reached a value that was steady over multiple windows.

The sigma-r plots for all constructs were made to show the average range of motion between alpha carbons as a function of interatomic distance for each 100-ns block of simulation (35). The average interatomic distances between alpha carbons were calculated for each pair of residues, along with the standard deviation. The standard deviations within intervals with the step size of Δ*r* = 0.5 ns were averaged to determine the average σ values for the interatomic distance interval using the equation

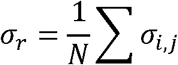

where *N* is the number of interatomic distances within the range

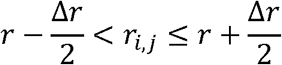

and *r*_*i,j*_, σ_*i,j*_ are the mean interatomic distance and standard deviation of the distance between alpha carbons *i* and *j* over 100-ns blocks of the construct trajectories. The values calculated for each 100-ns block were graphed and the standard deviations were calculated using the independent simulations.

To compare PC1 and PC2 for VCBC-CUL5 with each of the other constructs, we calculated the p-values for the mean and variance using a permutation test. The square of the difference between the mean of each independent simulation and the crystal structure was calculated and the variance within each independent simulation calculations were used to determine p-values. All p-values were found using 1000 permutations.

### Protein expression and purification

The VCBC complex, consisting of consensus Vif (19), CBF-β (residues 1-165), EloB (residues 1-118), and EloC (residues 17-112), was co-expressed in *E. coli* BL21 DE3 Star cells as described (19). Briefly, cells were grown to mid-log phase at 37 °C followed by induction with IPTG at 18 °C for 18 h. Cells were harvested and subsequently lysed by sonication in nickel buffer A (20 mM HEPES pH 8, 500 mM NaCl, 10% glycerol, 1 mM DTT, 20 mM imidazole), followed by centrifugation (14,500 g, 45 min, 4 °C) to clarify the lysate. The sample was purified by nickel chromatography, a heparin column, and size-exclusion chromatography into a pH 7.5 buffer containing 20 mM HEPES, 500 mM NaCl, 10% glycerol, and 1 mM DTT. To form the VCBC-dT14 complex, purified VCBC was concentrated and combined with oligonucleotide DNA (dT_14_) in a 1.5:1 ratio of protein to oligonucleotide, then diluted with 20 mM HEPES pH 7.5 to an NaCl concentration less than 40 mM, and purified by size exclusion chromatography in 20 mM HEPES, pH 7.5 buffer with 0.5 mM TCEP and 20 mM NaCl with no glycerol.

To express the VCBC complex with ^13^C isotope labels on the methyl groups (ILVMA) for NMR spectroscopy, the BL21 DE3 Star cells were grown in M9 minimal media with trace metals (36). Before inducing with IPTG, ^13^C isotope labeled amino acid precursors were added to the culture [50 mg/L 2-ketobutyric adid-4-^13^C, 100 mg/L 2-keto-3 methyl-^13^C-butyric acid-4-^13^C, 250 mg/L *L*-Methionine-methyl-^13^C1, 100 mg/L *L*-Alanine (3-^13^C)].

A solubility-optimized form of the A3F-C-terminal domain with 11 mutations (A3Fctd-11x) was expressed in *E. coli* BL21 DE3 Star cells as described (19, 37). The sample was purified by nickel chromatography and the His-tag was cleaved with TEV then passed again over the nickel column to remove the tag. The A3Fctd sample was then further purified by size-exclusion chromatography with a HiLoad Superdex 75 pg column (GE Healthcare). Budding yeast Upf1 was expressed and purified as described (38).

### Gel-shift assays

Poly(U)40 oligonucleotide (Dharmacon) was end-labeled by polynucleotide kinase with [γ-^32^P] ATP, according to the manufacturer’s recommendations (New England Biolabs). Reaction products were purified with an Illustra G-50 column (GE Healthcare). 100 pmol probe RNA was mixed with buffer or varying amounts of protein in NEB Buffer 2 and incubated at room temperature for 15 minutes. Native gel electrophoresis followed by phosphorimaging (Typhoon, GE Healthcare) was used to visualize the formation of RNA-protein complexes.

### RNAse protection

Body-labeled poly(U) RNA was synthesized by a tailing reaction with poly(U) polymerase (NEB), poly(U)_15_ oligonucleotide primer, and [α-^32^P] UTP. Long RNAs were purified using an RNeasy Mini Spin Column (QIAGEN) according to the manufacturer’s recommendations. For each RNAse protection assay, 1 μL of 2×10^5^ cpm / μL body-labeled poly(U) RNA was mixed with either 2 μM protein or buffer in 1/2 X NEB Buffer 2. RNAse A was added at a final concentration of between 2-10 μg/mL, and reactions were mixed and allowed to proceed for one hour at room temperature. Nuclease-resistant products were purified by two successive chloroform-phenol extractions followed by ethanol precipitation overnight at −80° C. Inclusion of glycogen at a final concentration of 15 μg/mL during ethanol precipitation was critical for the recovery of small RNA fragments. Finally, reaction products were visualized by denaturing gel electrophoresis and phosphorimaging (Typhoon, GE Healthcare) followed by analysis in ImageJ (39).

### Fluorescence anisotropy

5’ fluorescein-labeled and unlabeled oligonucleotides were purchased from IDT. All measurements were made using an Analyst AD system (LJL BioSystems). For direct binding assays, VCBC was serially diluted into a solution containing 25 mM HEPES pH 7.9, 25 mM NaCl, 1.25 mM MgCl2, 1 mM DTT, and 1 nM fluorescent probe. For competitive binding assays, an unlabeled competitor oligonucleotide was diluted into a solution containing 25 mM HEPES pH 7.9, 25 mM NaCl, 1.25 mM MgCl2, 1 mM DTT, 0.5 nM fluorescent probe, and VCBC at a concentration equal to the *K*_*d*_ value determined for probe-VCBC binding. All data was fit in R using the *minpack.lm* package (40); direct binding experiments were fit using a single-site saturation binding model, and competitive binding experiments were fit as described (41). Plots were generated using the ggplot2 package for R (42).

### Methyl-TROSY NMR spectroscopy

NMR was performed in 20 mM HEPES, pH 7.5, D_2_O buffer with 20 mM NaCl and 0.5 mM TCEP. NMR experiments were performed on an 800 MHz Bruker spectrometer with a cryogenically cooled probe. Methyl-TROSY experiments were performed at 300 K (43). The VCBC concentration was ∼160 μM for the VCBC-dT14 experiment and 256 scans were performed in the ^1^H dimension. The A3Fctd concentration was ∼108 μM and the VCBC concentration was ∼90 μM for the VCBC-dT14-A3Fctd experiment, and to adjust for this concentration difference, 816 scans were performed in the ^1^H dimension. Spectra were processed using the NMRPipe suite of programs (44). Figures and chemical shift assignments were done using the python module nmrglue (45). Remaining data analysis was performed using in-house python scripts.

### Isothermal titration calorimetry

The binding affinity between A3Fctd and VCBC was measured using a VP-ITC Micro Calorimeter (MicroCal Inc.). The proteins A3F and VCBC were dialyzed extensively against either high salt buffer 20 mM Hepes 7.5, 10% Glycerol, 300 mM NaCl (A3F into VCBC monomer) or low salt buffer 20 mM Hepes 7.5, 10% Glycerol, 50 mM NaCl, 2 mM dT14 (A3F into VCBC dimer) prior to titration. The injection syringe and the sample cell were filled with A3F and VCBC, respectively. The system was equilibrated to a stable baseline before initiating an automated titration. Each titration experiment injected 7 μL of A3F and 30 injections were repeated at 200 s intervals at 25°C. The sample cell was stirred at 300 rpm. Heat changes upon the addition of the A3F were monitored and fitted using the least-squares regression analysis of ORIGIN software (MicroCal Inc.) to calculate the dissociation constant (K_d_).

### Size-exclusion chromatography multi-angle laser light scattering

To determine the molecular size of VCBC momomer and dimer, the purified proteins were injected into Shodex KW-804 size exclusion column with an Ettan LC (GE Healthcare) and the elution products were applied to inline DAWN HELEOS MALS and Optilab rEX differential refractive index detectors (Wyatt Technology Corporation). Data were analyzed by the ASTRA V software package (Wyatt Technology Corporation).

## Results

### Alternate conformations of the VCBC complex

#### Simulations of VCBC and VCBC-CUL5ntd

We performed MD simulations of the VCBC complex alone or bound to CUL5ntd. In both simulations we observed large-scale global motions of the complex correlated with flexibility in the Vif linker region, near the Zn^2+^ coordination site. We used Cartesian coordinate PCA to identify the global motions that were most dominant in our simulations. To compare the VCBC and VCBC-CUL5ntd simulations, we performed PCA on the combined ensemble all of the VCBC structures from the last 300 ns of both the VCBC and VCBC-CUL5ntd simulations. The first two PCs accounted for ∼71% of the variance in the VCBC structure over both sets of simulations (with and without CUL5ntd). The first PC (48% of the variance) involved a twisting open of the Vif α/β domain and CBF-β with respect to the Vif α-domain and EloBC, while the second PC (23% of the variance) captured a clamshell motion where the Vif α/β domain and CBF-β come closer to EloBC (Fig. 2). Figure 2 shows that while the simulations with CUL5ntd do include a small amount of sampling along these PCs, VCBC in the absence of CUL5 samples conformations substantially different from the crystal structure, indicating that VCBC adopts alternate conformational states not present when CUL5ntd is bound. Table 1 shows the mean and standard deviation for PC1 (twisting motion) and PC2 (clamshell motion) for each construct. When we performed PCA on only the VCBC-CUL5ntd ensemble, there was a large overlap (> 50%) of each of first two PCs with those shown in Fig. 2 for the VCBC/VCBC-CUL5ntd combined ensemble (Table S3 in the Supporting Material). This indicates that the same types of motions are occurring for VCBC-CUL5ntd, but the total conformational change is not as large.

**Figure 2.**
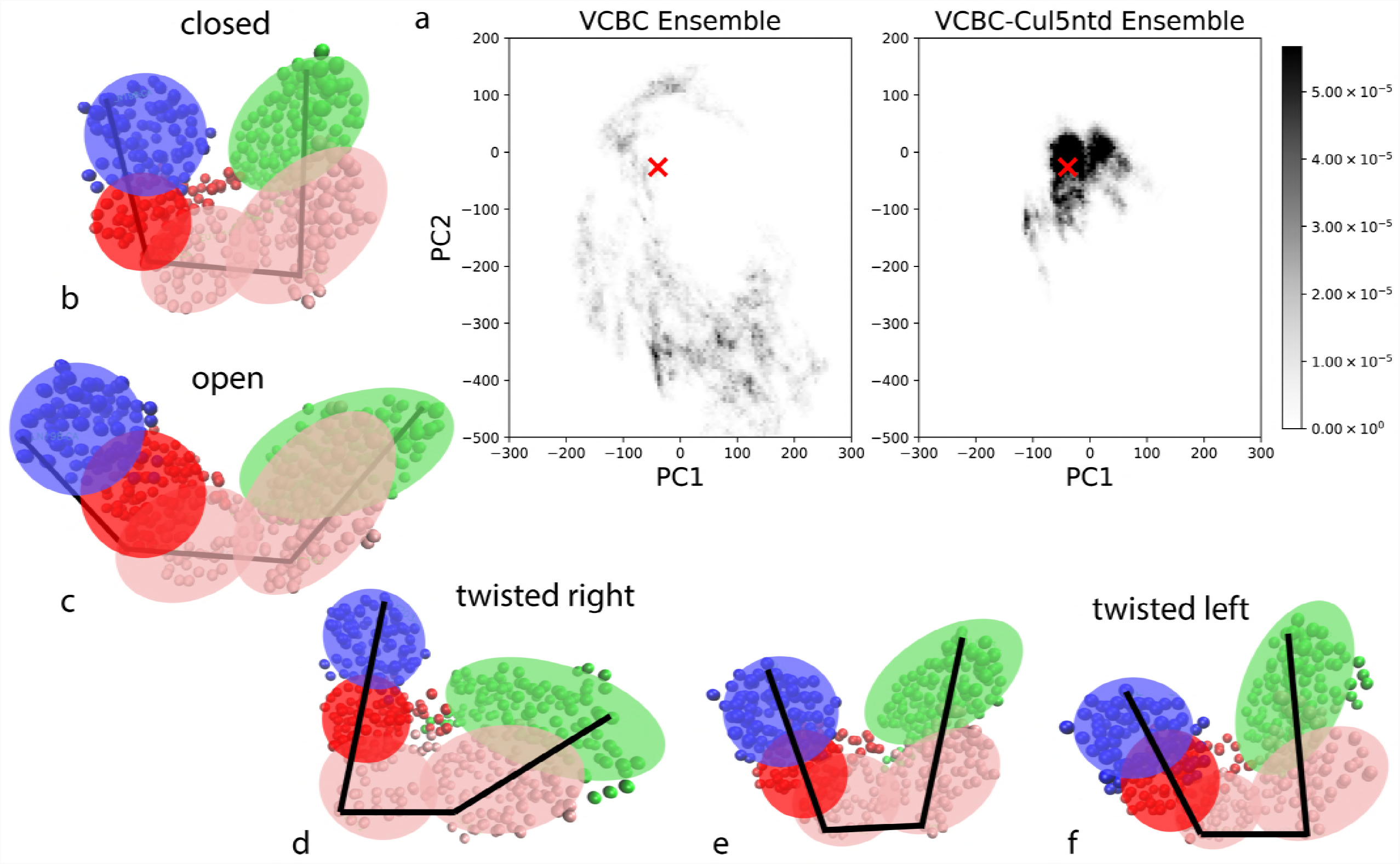
a) Projection of the VCBC and VCBC-CUL5ntd ensembles onto the first two principal components of motion. The shading of the cell represents the fraction of the ensemble at that value of PC1 and PC2, where black is 5.4 x 10^−5^ or greater and white is 0. The red x represents the values of PC1 and PC2 in the crystal structure from which all simulations were initiated. d-f) PC1 represented as a series of Cα positions in the range plotted with a cartoon overlaid. Vif is shown in pink, CBF-β in green, EloB in blue, and EloC in red. b,c) PC2 represented as Cα positions in the range plotted with a cartoon overlaid.

**Table 1.**
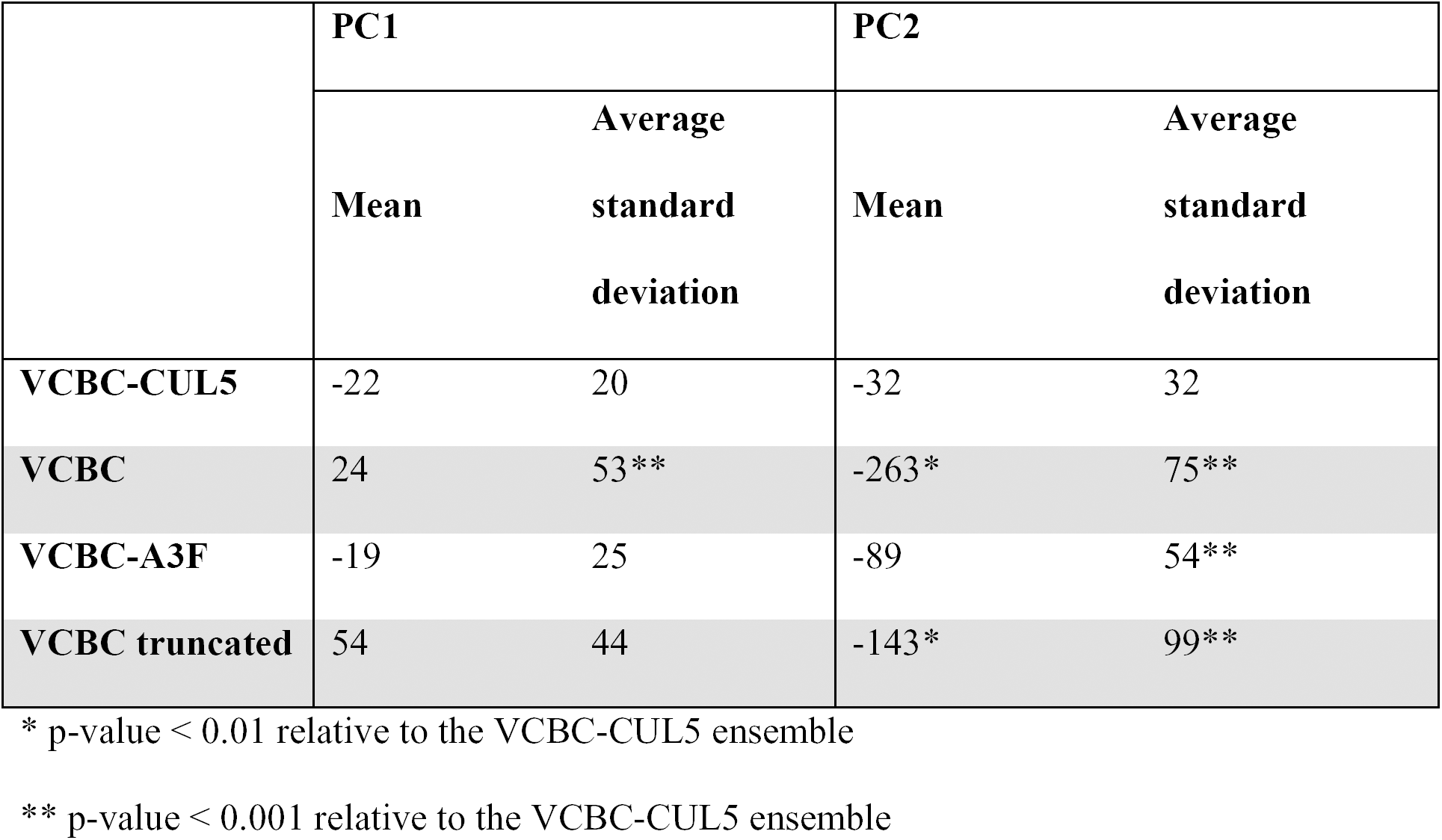
Comparison of average PC value for each construct simulated.

|  | **PC1** |  | **PC2** |  |
| --- | --- | --- | --- | --- |
|  | **Mean** | **Average**<br>**standard**<br>**deviation** | **Mean** | **Average**<br>**standard**<br>**deviation** |
| **VCBC-CUL5** | -22 | 20 | -32 | 32 |
| **VCBC** | 24 | 53** | -263* | 75** |
| **VCBC-A3F** | -19 | 25 | -89 | 54** |
| **VCBC truncated** | 54 | 44 | -143* | 99** |

\* p-value < 0.01 relative to the VCBC-CUL5 ensemble
\*\* p-value < 0.001 relative to the VCBC-CUL5 ensemble

We measured the autocorrelation time for each of the first two PCs in order to determine the time scale of the motions that these PCs are capturing. VCBC exhibited autocorrelation times of ∼25 ns for both PC1 and PC2, meaning that (at least) hundreds of nanoseconds of simulation time would be required to fully sample those motions. To make sure that we had run each of the individual simulations long enough for the dynamics of the complex not to be affected by the starting crystal structure conformation, we compared the global dynamics of the VCBC complex over sequential 100 ns blocks of simulation time using sigma-r plots (35) (Fig. S2 in the Supporting Material). The similarity of the global motions based on sigma-r plots, as well as the magnitude of the first five eigenvalues from PCA (Fig. S1) show that the VCBC simulations had equilibrated after 300 ns and the VCBC-CUL5ntd simulations after 100 ns. We collected 300 ns of simulation time after this equilibration period from eight independent simulations to compare the VCBC and VCBC-CUL5ntd ensembles.

To investigate local conformational changes associated with the global dynamics of the VCBC complex observed by PCA, we measured protein backbone conformational changes and hydrogen bonding patterns. Overall, there were no major conformational changes within each of the protein domains, and most secondary structure elements were maintained to a similar degree in the VCBC and VCBC-CUL5ntd simulations. The main region of the complex that showed local conformational differences between the two ensembles was the Vif linker region, between the α-domain (that interacts with EloC and CUL5) and α/β-domain (that interacts with CBF-β). We observed a small increase in root-mean-squared fluctuations of the Vif linker backbone when CUL5 was not present (Fig. S4 in the Supporting Material). Additionally, α-helices adjacent to the linker were slightly destabilized in the VCBC simulations without CUL5ntd. Rearrangement of this linker region is likely necessary for the global conformational changes that reorient the other proteins relative to each other, and this results in greater flexibility in this region when there are global conformational changes.

We also examined the backbone dihedral angles sampled by residues in the Vif linker region (residues 25-30, 111-117, and 126-173). We identified several residues that can sample multiple backbone conformations. Many of these residues, shown as spheres in Figure 3c, sample a different distribution of backbone dihedrals when VCBC is not bound to CUL5, even though they do not directly interact with CUL5. These alternate dihedral conformations are associated with the alternate conformations of the complex from PCA (Fig. 3). Flexible dihedral angles in this Vif linker region are likely necessary for the overall expansion, contraction, and twisting of the VCBC complex that occurs when CUL5 is not bound.

**Figure 3.**
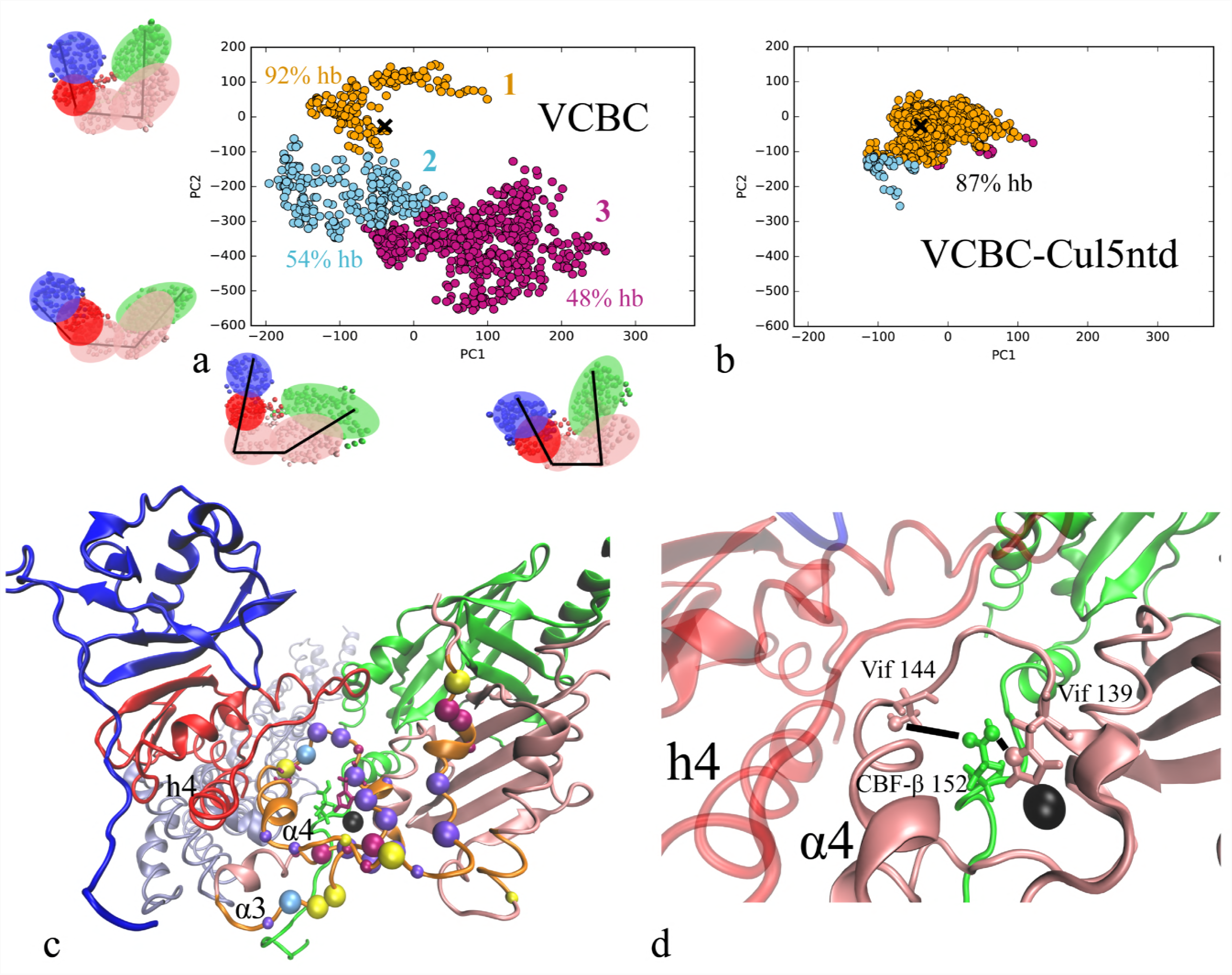
Alternate conformations of VCBC. a) Projection of the VCBC ensemble onto the first two principal components of motion. Clustering was performed based on PC1 and PC2, and the color represents the membership in three different clusters. The clusters are labeled with the percent of structures in that cluster where at least *one* of the two Vif linker hydrogen bonds with CBF-β 152 are formed (Vif 144 or Vif 139). b) Projection of the VCBC-CUL5ntd ensemble onto the first two principal components of motion. The color of the data points indicates which of the three clusters from the VCBC clustering (plot a) the VCBC-CUL5ntd data is classified into. The black x in plots a) and b) represents the values of PC1 and PC2 in the crystal structure from which all simulations were initiated. c) VCBC structure (Vif is shown in pink, CBF-β in green, EloB in blue, and EloC in red, zinc ion in black) with Vif region where dihedrals were measured shown in orange. Residues that showed a significant difference in the mean phi or psi angle (or both) between the VCBC-CUL5ntd simulations and VCBC simulations are shown as spheres. The large spheres indicate residues that had a difference of the mean angle of more than 20° between the VCBC-CUL5ntd and VCBC simulations. Magenta spheres indicate backbone dihedral angles that are different from the VCBC-CUL5ntd only in cluster 3 of the VCBC ensemble; light blue spheres indicate backbone dihedral angles that are different from the VCBC-CUL5ntd dihedral angle only in cluster 2 of the VCBC ensemble; and purple spheres indicate backbone dihedral angles that are different from the VCBC-CUL5ntd dihedral angle in clusters 2 and 3 of the VCBC ensemble, but not in cluster 1. Yellow spheres differ from the VCBC-CUL5ntd dihedral angle in cluster 1 of the VCBC ensemble. Vif residues 144 and 139 sidechains are shown in pink because these residues form hydrogen bonds with CBF-β 152 (sidechain shown in green) mostly in cluster 1 (like in the VCBC-CUL5ntd ensemble) and not in cluster 3. Vif helices α3 and α4 and EloC helix h4 are labeled. d) Close up of sidechains involved in linker hydrogen bonds, with black lines indicating where the hydrogen bonds form.

A comparison of hydrogen bonding patterns also points toward the Vif linker region. We compared all of the hydrogen bonds present in the complex and identified which residues show a difference in hydrogen bonding pattern in the VCBC simulations compared to the VCBC-CUL5ntd simulations (Fig. S6 in the Supporting Material). The only hydrogen bonds that showed a large difference between the two ensembles are shown in Figure 3d and are in the Vif linker region. These residues, His139 (part of the HCCH zinc binding motif) and Ser144, hydrogen bond with Glu152 on the CBF-β C-terminal tail much more often in the VCBC-CUL5ntd simulations than in the simulations with only VCBC, though they do not interact directly with CUL5. In cluster 1 of the VCBC ensemble, 92% of structures contain at least one of these two hydrogen bonds, while in about half of the structures in clusters 2 and 3 neither hydrogen bond is present. These hydrogen bonds may stabilize the cluster 1 VCBC conformation that is dominant when bound to CUL5ntd. Overall, the flexibility of the Vif linker region is correlated with the large-scale dynamics we observe in our MD simulations.

### NMR of the VCBC complex

To experimentally observe the conformational heterogeneity of the VCBC complex, we performed methyl-TROSY NMR spectroscopy on ILVMA ^13^C-labeled VCBC. Because only ILVMA residues are visible, the number of cross-peaks in the 2D spectrum are reduced, and therefore, this technique enables observations of methyl resonances in molecular assemblies as large as the proteasome (46). The resonance intensities are sensitive to dephasing that occurs due to conformational changes on the μs-ms timescale, including global conformational changes or self-association, which, in the limit of intermediate exchange, would cause peaks to broaden and disappear (47). The VCBC complex is difficult to work with at the concentrations needed for NMR spectroscopy due to its tendency to aggregate and precipitate at high concentration. To prevent aggregation, the complex can be kept in a buffer with high salt (500 mM NaCl) and 10% glycerol, but the NMR signal is broadened under these conditions and individual methyl peaks are not visible. We found that a defined length of single-stranded DNA will bind the VCBC complex (Fig. S7 in the Supporting Material) and enable it to be concentrated under low-salt conditions (20 mM NaCl) with no glycerol present, possibly by shielding the positively charged Vif from non-specific electrostatic interactions with other copies of the complex. Using fluorescence anisotropy measurements, we found that the optimal single-stranded DNA length is 14 oligonucleotides (dT14), binding to VCBC with low nanomolar affinity, *K*_*d*_ = 3.7 ± 1.7 nM (Fig. 4a and b). We also noticed that VCBC forms a dimer when DNA is present, as observed by size-exclusion chromatography (SEC) in line with multi-angle laser light scattering (MALLS) (Fig. 4c). Since many CRL complexes are known to form dimers, and the dimerization is often mediated by the substrate binding or adaptor proteins, this dimer of the Vif complex may be the functionally relevant species (48–55).

**Figure 4.**
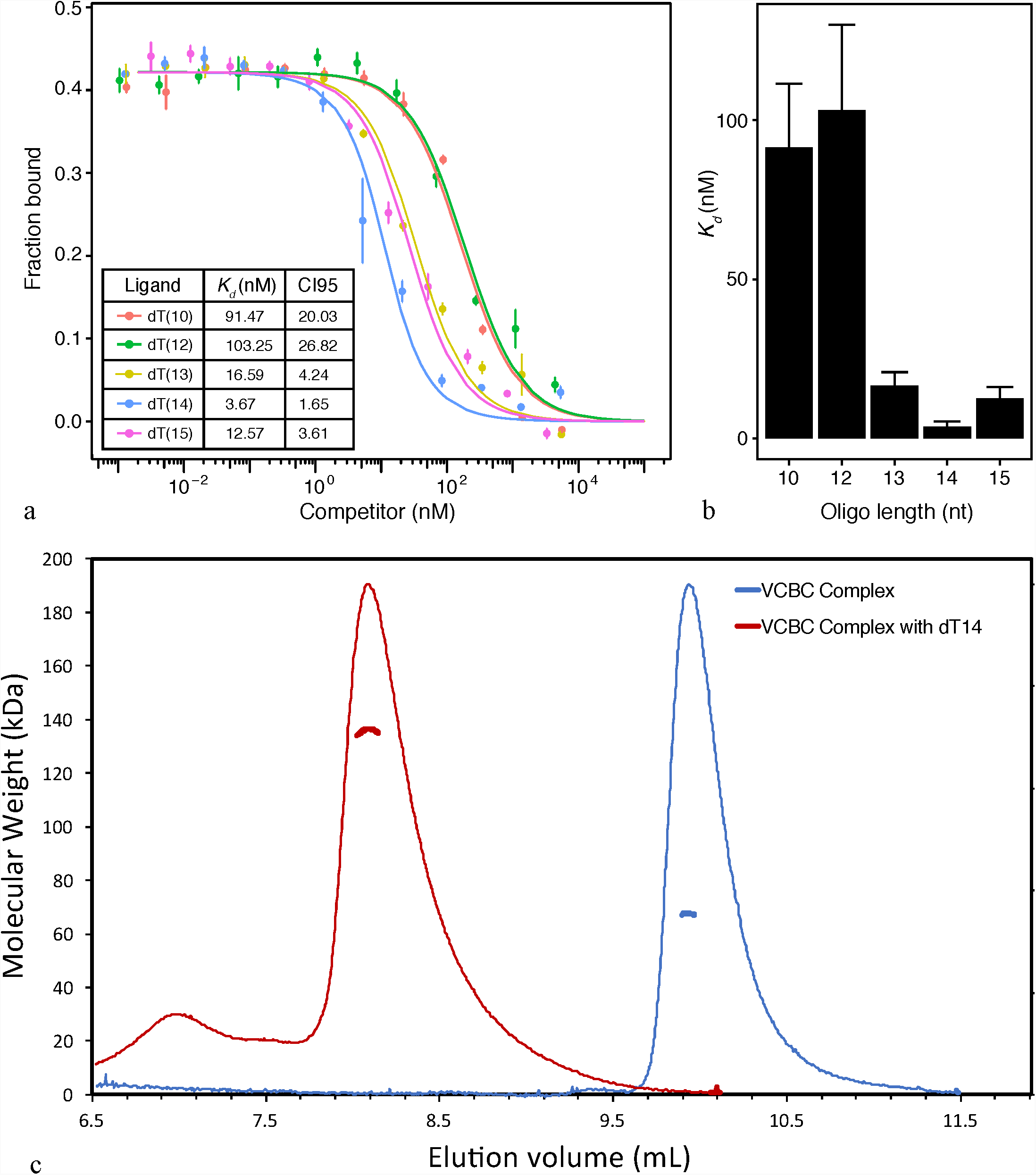
Characterization of VCBC interactions with DNA. a) Fluorescence anisotropy data for VCBC-oligonucleotide interactions under competitive binding conditions. Points are the means of three independent experiments, and error bars indicate the standard deviation. b) K_d_ values determined for the fitted curves shown in panel a. Error bars indicate CI95. c) Size Exclusion Chromatography with Multi-Angle Laser Light Scattering (SEC-MALLS) of the VCBC complex in the absence (blue) and presence (red) of dT14. The elution volume of the VCBC complex in 300 mM NaCl (blue) corresponds to an experimental molecular mass of 67 kDa, which is consistent with the theoretical molecular weight for a monomer of 66 kDa. The elution volume of VCBC complex bound to dT14 in 50 mM NaCl (red) corresponds to an experimental molecular mass of 135 kDa, which is consistent with the theoretical molecular weight for a monomer of 66 kDa, indicating that the complex is a dimer.

We collected methyl-TROSY NMR of the ^13^C-ILVMA-labeled VCBC complex bound to dT15 (Fig. 5). The resulting VCBC-dT14 spectrum contains few observable methyl peaks. For example, out of 28 Ile residues in the VCBC complex, only 2 or 3 peaks in the Ile-C_δ_ region of the spectrum were resolved. The absence of expected methyl peaks indicates that dynamics on the μs-ms timescale are causing peak broadening due to motions occurring on the NMR chemical shift timescale (i.e., the intermediate exchange regime). While the timescale is longer than the ns dynamics observed in our simulations, the broadening of most methyl peaks is consistent with the large-scale conformational rearrangements from MD. Table S4 in the Supporting Material gives the number of methyl groups that are present in the complex, of which only a small number are visible.

**Figure 5.**
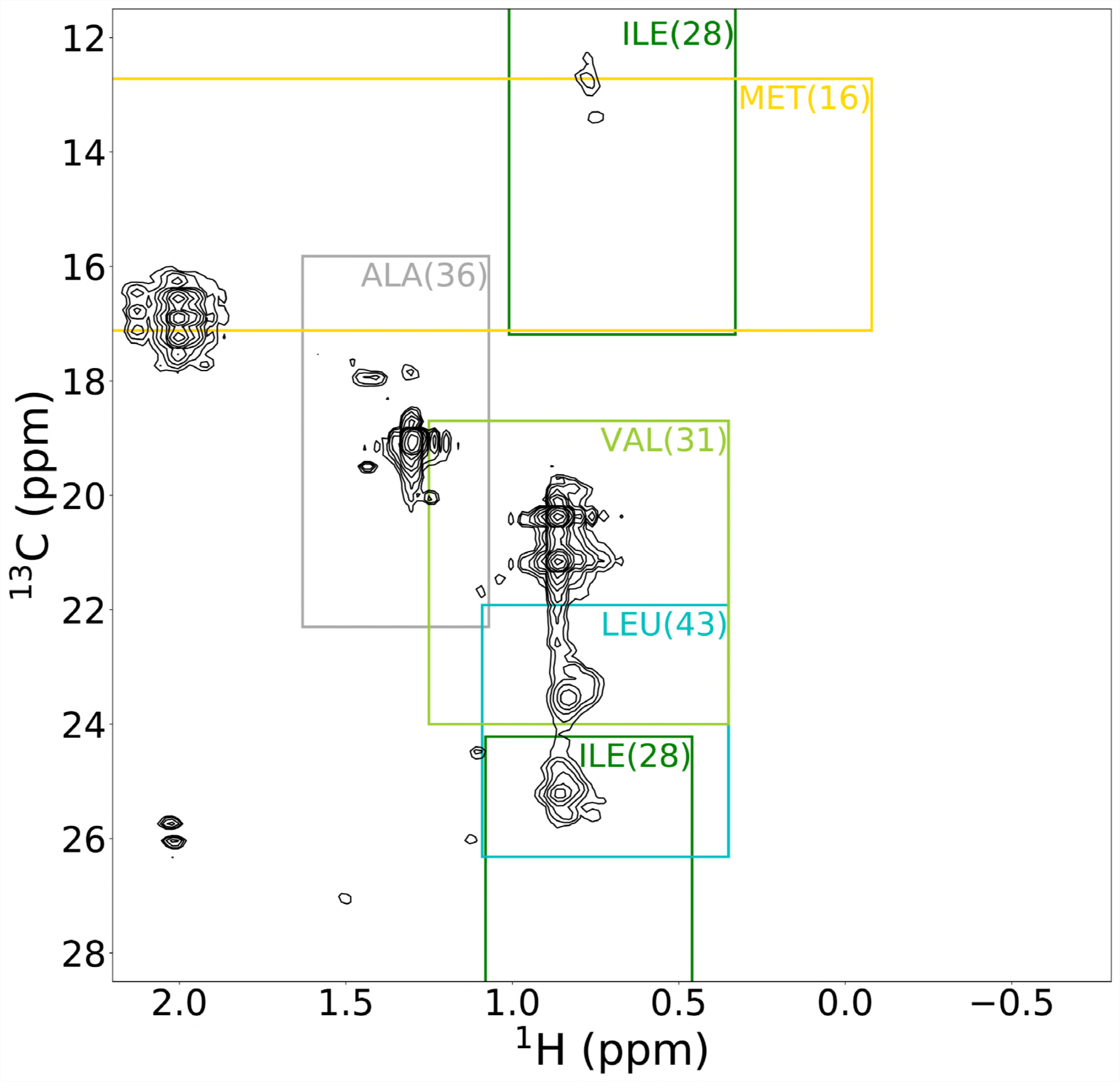
^13^C-^1^H Methyl-TROSY spectrum of the VCBC complex with ^13^C isotope labels on Ile, Leu, Val, Met, Ala residues recorded at 800 MHz at 300 K. 14-T ss-DNA is present to stabilize the VCBC complex. Boxes indicate the typical range where the methyl peaks from each residue type are found **(56)**. The number in parentheses indicates the number of peaks we would expect to see in that region given the sequences of the proteins present in the VCBC complex.

### Effect of A3Fctd on the dynamics of the VCBC complex

#### NMR of VCBC-A3Fctd

To further test predictions of the MD simulations, we hoped to collect NMR data of the VCBC complex with an additional protein partner bound to observe any change in the conformational exchange visible in the methyl-TROSY data. We attempted these experiments with the CUL5ntd bound as in our simulations, however these NMR experiments were not tractable and could not yield resolved spectra even with deuterated VCBC proteins. While we were not able to collect NMR data on the VCBC-CUL5 complex, other proteins that bind the VCBC complex could be hypothesized to have similar effects on the overall dynamics of the complex, and therefore we decided to investigate the VCBC complex bound to a substrate protein, specifically A3F. Before performing NMR experiments, we wanted to ensure that the presence of DNA would not influence the way that A3F interacts with the VCBC complex. Isothermal titration calorimetry (ITC) experiments showed that A3Fctd, which is the domain that binds to Vif (17), binds to VCBC (in high salt) and VCBC-dT14 (in low salt) with nearly identical affinities (Fig. S9 in the Supporting Material). ITC experiments show that the dimerization of the VCBC-dT14 complex does not affect VCBC affinity for A3F, and additional SEC experiments show that VCBC-dT14 remains a dimer even after A3F binds (Fig. S10 in the Supporting Material).

The VCBC-dT14-A3Fctd construct had better spectroscopic behavior than VCBC-dT14-CUL5ntd, and we were able to resolve a methyl-TROSY spectrum of VCBC from this construct (Fig. 6, red spectrum), which contained many more visible methyl-TROSY peaks than the VCBC spectrum without A3Fctd bound (Fig. 6, Table S4). In total, we observed at least 50 distinct additional methyl resonances for VCBC with A3Fctd bound compared to without A3Fctd. While not all of the methyl groups in the VCBC complex were resolved for the VCBC complex bound to A3F, substantially more peaks were present, indicating that binding of A3Fctd reduces the conformational exchange on the intermediate timescale for several ILVMA residues in the complex, resulting in less broadening of those resonances. For example, out of 28 Ile residues in the VCBC complex, we see about 15 peaks in the Ile-_δ_ region of the spectrum for the VCBC complex bound to A3Fctd, rather than only 2 or 3 peaks for the VCBC complex without A3Fctd. A3Fctd binding must be the cause of the reduction in dynamics, since both experiments were performed on VCBC with dT14 present. As in our simulations with and without CUL5 bound, this indicates that the VCBC complex is dynamic and occupies alternate conformational states which are restricted when another protein partner binds to the complex and alters the conformational energy landscape.

**Figure 6.**
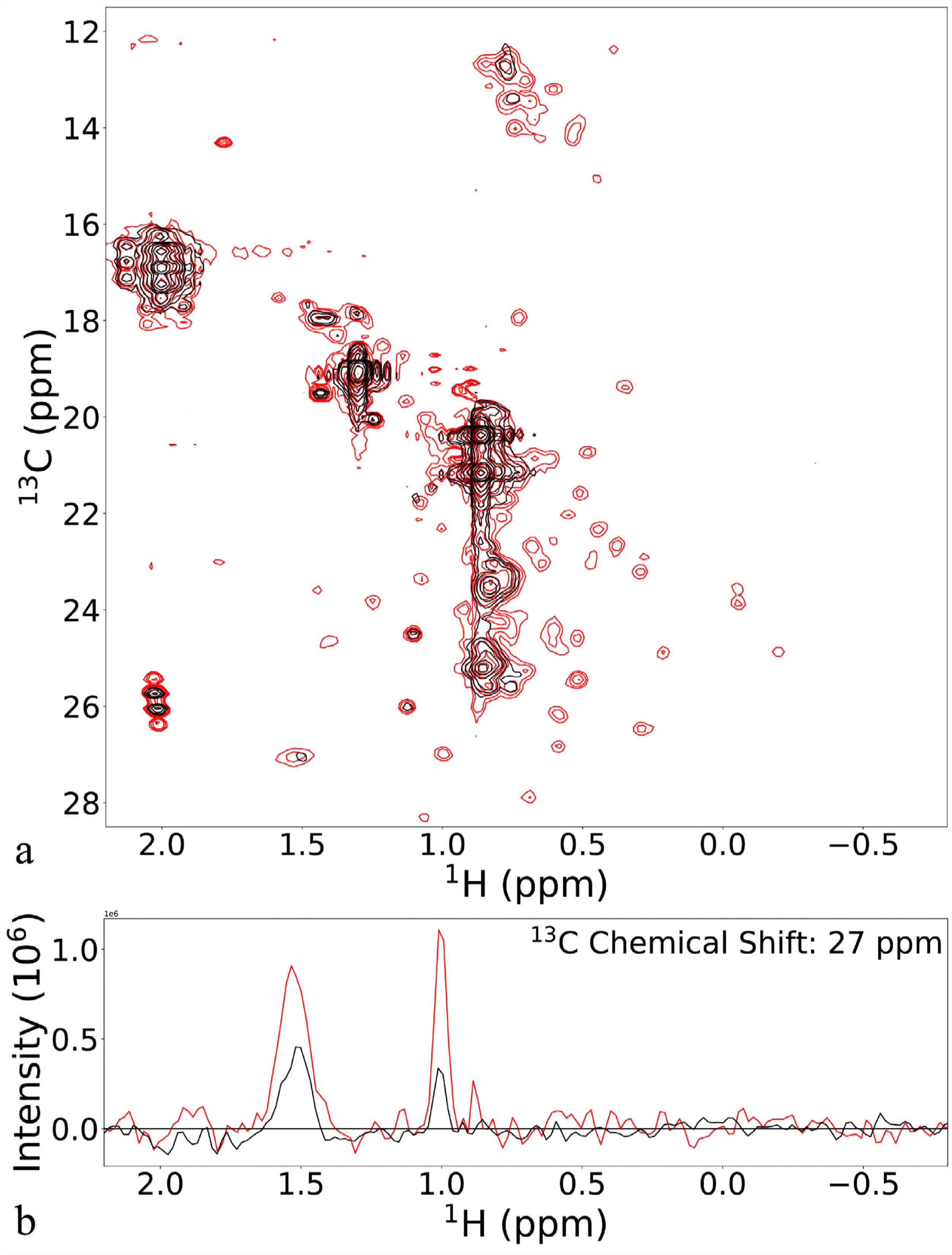
^13^C-^1^H Methyl-TROSY spectrum of the VCBC complex with and without A3Fctd. a) ^13^C-^1^H Methyl-TROSY spectrum of the VCBC complex bound to the A3Fctd (red) with the spectrum of the VCBC complex (black). The VCBC complex proteins have ^13^C isotope labels on Ile, Leu, Val, Met, Ala residues, while the A3Fctd is unlabeled. Both spectra have 14-T ss-DNA present to stabilize the VCBC complex and are recorded at 800 MHz at 300 K. b) One-dimensional cross-section of the spectrum at ^13^C chemical shift of 27 ppm showing two of the methyl peaks.

#### MD simulations of the VCBC complex bound to A3Fctd

Based on our simulations of VCBC-CUL5ntd and the Methyl-TROSY data of the VCBC complex bound to A3Fctd, we suspected that A3Fctd might have a similar effect to CUL5ntd, restricting the global motions of the VCBC complex. There is no solved experimental structure of A3Fctd bound to Vif that could be used to initiate MD simulations of the VCBC-A3Fctd complex, but Richards *et al.* have published a modeled structure of the Vif-A3F interface based on mutational data (26). We used this model to initiate MD simulations of the VCBC-A3Fctd complex. After running four independent simulations of the VCBC-A3Fctd complex for 400 ns, we projected the conformations sampled in the last 300 ns of each simulation onto the coordinates of correlated motion that were identified based on PCA of the VCBC and VCBC-CUL5 simulations to determine if the complex exhibited similar motions when bound to A3Fctd (Fig. 7). The mean and standard deviations for PC1 and PC2 are given in Table 1. While the VCBC complex bound to A3F samples a wider range of values than the VCBC-CUL5 complex along PC2, it does not sample as many alternate conformations as the VCBC complex, and generally adopts a conformation that is closer to the VCBC-CUL5ntd crystal structure. Consistent with our NMR data (Fig. 6), the comparison of VCBC and VCBC-A3F in terms of these global conformational changes indicates that the binding of A3F reduces the dynamics of the VCBC complex.

**Figure 7.**
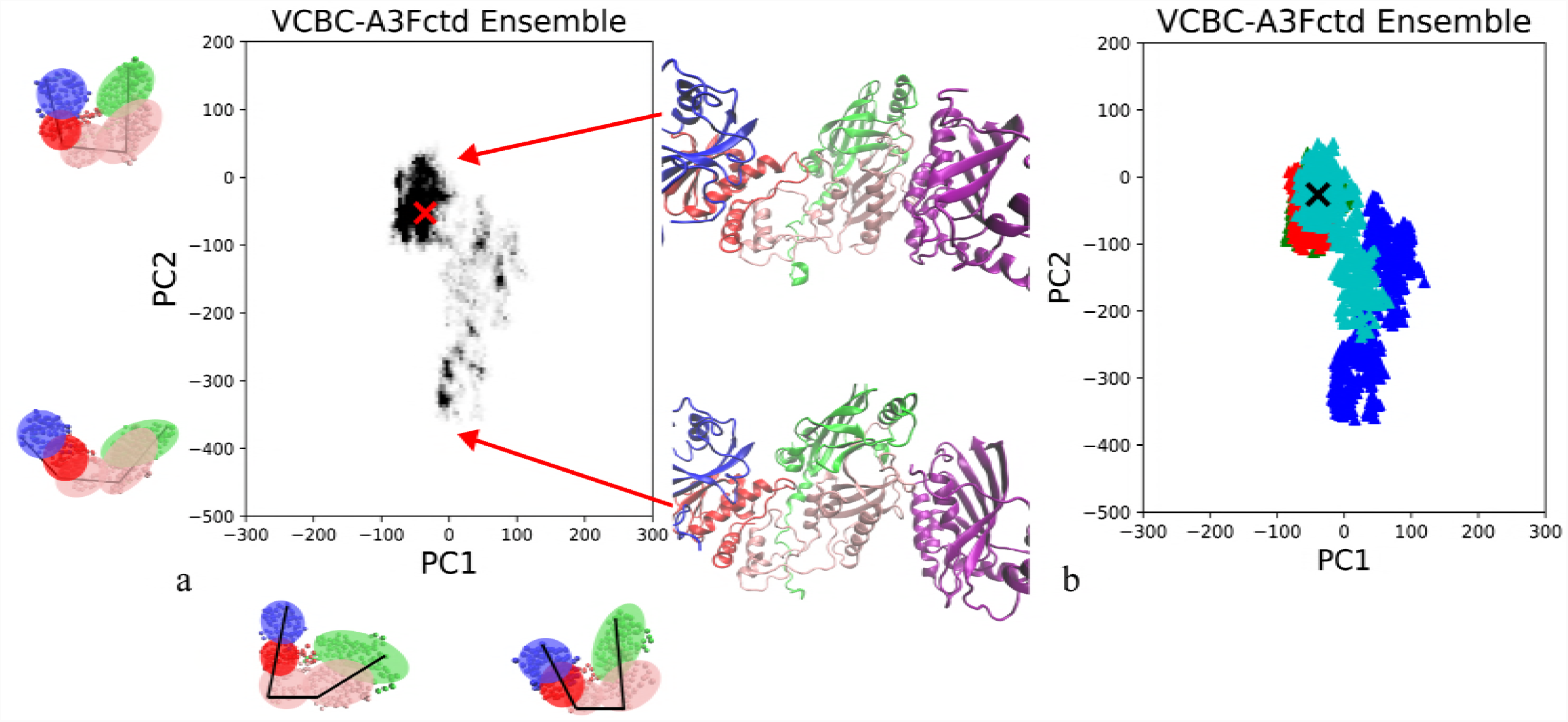
Projection of the VCBC-A3Fctd ensembles onto the first two principal components of motion. In (a), the shading of the cell represents the fraction of the ensemble at that value of PC1 and PC2, where black is 5.4 x 10^−5^ or greater and white is 0. In (b) each independent simulation is shown in a different color. The x represents the values of PC1 and PC2 in the crystal structure from which all simulations were initiated.

The VCBC-A3Fctd simulations were not as consistent as the VCBC or VCBC-CUL5ntd simulations. As shown in Figure 7, two of the four VCBC-A3Fctd simulations sampled states close to the crystal structure (similarly to VCBC-CUL5ntd), while two simulations sampled more widely along PC 2, indicating a clamshell opening of the complex. The VCBC-A3Fctd trajectories with a less stable conformation may have resulted from an initial unstable A3F-Vif interface, since this construct was initiated based on a computational model and not on an experimentally determined structure. The two simulations that diverged the most from the crystal structure along PC1 and PC2 also showed some separation of the A3Fctd protein from the rest of the VCBC complex (Fig. 7). Therefore, the increased dynamics of VCBC in complex in these simulations may occur because A3F begins to unbind from the rest of the complex due to a high-energy initial structure.

## Discussion

All-atom simulations of the VCBC complex predict that this complex is conformationally heterogenous, exhibiting large twisting and clamshell motions that involve a flexible hinge in the Vif linker region near the zinc coordination site. By contrast, the crystal structure of the VCBC-CUL5ntd complex shows a well-defined and compact tertiary structure (4). However, several types of experimental data support increased conformational heterogeneity of the VCBC complex as observed in our MD simulations. Our NMR studies support the model that VCBC undergoes global conformational changes on the μs-ms timescale, and that binding of the A3Fctd protein, and likely other proteins such as CUL5 or additional A3 proteins, to the complex reduces these motions allosterically. While our MD simulations indicate that the VCBC-A3Fctd retains flexibility, this may be affected by the starting structure of VCBC-A3F, which is based on computational modeling (17), since the binding interface has not been solved from structural experiments.

Previously reported SAXS data also indicates that the VCBC complex may be more flexible alone than when bound to CUL5 (6). The envelope calculated by SAXS was more elongated than the VCBC complex bound to CUL5 in the crystal structure, indicating that the VCBC complex alone can sample more extended conformations. A recent electron microscopy (EM) study of VCBC bound to antibody antigen-binding fragments (Fabs) also indicates flexibility in the VCBC complex, likely caused by the hinge at the linker region between the two Vif domains (19). The magnitude of conformation heterogeneity of VCBC we observe by MD and NMR would likely make EM of the VCBC complex alone intractable, but the binding of the Fabs stabilizes the complex enough that Binning *et al.* were able to classify the EM micrographs into two structurally similar EM maps (19). Overall, data from experiments and our simulations indicate that while the HIV-1 Vif protein can fold and form a stable complex with only EloB, EloC and CBF-β, additional binding partners further restrict Vif’s conformational landscape.

Previous MD simulations on a human CRL complex by Liu and Nussinov revealed conformational changes within the linker between the two domains of the substrate binding protein that correlated with changes in distance between the E2 ligase and substrate, which are critical for ubiquitin transfer (12, 13). Similarly, in our VCBC simulations, the flexibility of the Vif linker region facilitates conformational changes that could assist in the accurate positioning of APOBEC3 proteins for ubiquitination. Notably, while VCBC exhibits larger conformational changes in our simulations, the VCBC-CUL5 complex also undergoes the same type of global motions involving rotation around the hinge in the Vif linker but to a smaller degree. These conformational changes may therefore take place within the Vif CRL holoenzyme complex during the ubiquitination process.

Conformational flexibility of the VCBC complex may also be relevant for its ability to ubiquitinate multiple APOBEC3 proteins, some of which, like A3F and A3G, bind the ubiquitination complex at different sites on Vif (10, 11, 16, 57). While it is possible that all APOBEC3 proteins could rigidify the VCBC complex to the same extent and in the same conformation, other intrinsically disordered proteins have been demonstrated to adopt different conformations with their multiple binding partners (58, 59). Recent work by Binning *et al.* also showed that Vif disrupts the APOBEC3 packaging into virions by a degradation-independent mechanism that occurs when CUL5 is prevented from binding to the VCBC complex (19). The mechanism by which Vif prevents APOBEC3 packaging is independent of CUL5 binding and could potentially involve alternate conformations of the VCBC complex as observed in our molecular dynamics simulations. Much remains to be learned about how Vif functions in human cells, and the conformational heterogeneity that it exhibits in complex with human proteins could be important for enabling its multiple functions.

Finally, alternate conformations of the VCBC complex, whether or not they are functionally relevant, may provide a strategy for therapeutically targeting Vif. A small molecule that could trap Vif in a nonfunctional conformation, e.g., by binding selectively to certain conformations of the linker region, could prevent APOBEC3 ubiquitination and degradation, or potentially prevent APOBEC3 from binding VCBC at all. Allosterically blocking CUL5 binding to VCBC is also theoretically possible, but APOBEC3 is still prevented from packaging when CUL5 binding is blocked (19), as discussed above. Future studies of the conformational ensemble of VCBC with mutations in the Vif linker region would help to test the hypothesis that changes to the local interactions in this region could restrict the VCBC conformational ensemble, and could set the stage for therapeutic developments.

## Author Contributions

KAB proposed the project. KAB, JDG, JMB, and MPJ designed the research approach. KAB, LMC, ET, and ST performed the simulations and analyzed the data. KAB, DJS, HMT, and LAB performed the experiments and analyzed the data. KAB, LMC, JMB, MPJ, and JDG wrote the paper. KAB, LMC, DJS, ET, ST, HMT, LAB, JMB, MPJ, and JDG edited the paper.

## ACKNOWLEDGEMENTS

We thank Hiroshi Matsuo, National Cancer Institute, for the kind gift of A3F-CTD-11x plasmid; and Elena Conti of the Max Planck Institut für Biochemie for the gift of the budding yeast Upf1 expression plasmid. This work was supported by National Institutes of Health grant P50 GM082250 to MPJ and JDG; by F32 GM114894 to KAB; and F32 AI120867 to JMB, Support was provided by Skidmore College to KAB and by the Skidmore Faculty Student Summer Research Program to LMC and ET.

